# Differential effects on neuromuscular physiology between *Sod* loss-of-function mutation and paraquat-induced oxidative stress in *Drosophila*

**DOI:** 10.1101/2021.03.08.434419

**Authors:** Atsushi Ueda, Atulya Iyengar, Chun-Fang Wu

**Author notes:** **Corresponding author**: Dr. Atulya Iyengar, Department of Biology, University of Iowa, Iowa City, IA 52242. **Species**: *Drosophila melanogaster. **Data Type**: Phenotype Data. **Findings**: New Finding. **Author Contribution:** AU, AI and CFW designed the experiments. AU did the larval NMJ electrophysiology experiments, AI did the adult giant-fiber experiments. AI, AU and CFW analyzed data, and wrote the manuscript.

## Abstract

Oxidative stress is thought to be a major contributor to aging processes. Here, we report differential effects on neurotransmission caused by loss-of-function mutations of *Superoxide dismutase* (*Sod*) and by paraquat (PQ) feeding in *Drosophila*. We demonstrated alterations in *Sod* mutants; the larval neuromuscular junction displayed supernumerary discharges and the adult giant-fiber escape pathway showed increased latency and poor response to repetitive high-frequency stimulation. Even though the concentrations used led to motor coordination defects and lethality, PQ feeding failed to reproduce such performance deficits in these larval and adult preparations, indicating mechanistic distinctions between genetic and pharmacological manipulation of oxidative stress.

## Description

Free radicals, such as the superoxide anion (O_2_^−^) impart oxidative stress upon an animal and are thought to be a major contributing factor to age-related changes in the nervous system (Finkel & Holbrook, 2000; Harman, 1956; Harman, 1981). In eukaryotes, the cytosolic enzyme Cu^2+^/Zn^2+^ Superoxide Dismutase (encoded by *Sod* in *Drosophila*) is an important free radical scavenger that converts superoxide into hydrogen peroxide. *Drosophila Sod* loss-of-function mutants exhibit elevated levels of reactive oxygen species (ROS) combined with a drastically shortened adult lifespan (median ~11 d. vs 50 d. for wild-type flies at 25 °C, Phillips et al., 1989) and reduced locomotor ability in larvae (Şahin et al., 2017) and adults (Ruan & Wu, 2008). Coupled with these longevity and behavioral phenotypes, several neuromuscular deficits have recently been identified in *Sod* loss-of-function mutants including deranged nerve and synapse morphology in larvae (Milton et al., 2011) and adults (Agudelo et al., 2020; Şahin et al., 2017), as well as disrupted neurotransmission along the giant-fiber (GF) jump-and-flight escape circuit (Iyengar et al., 2020; Iyengar, 2016; Ruan, 2008). Given the striking phenotypes, a question that arises is: To what extent the straightforward assumption holds that aspects of the *Sod* phenotypes can be recapitulated by pharmacologically induced oxidative stress?

Here, we compare neuromuscular physiology in *Sod* loss-of-function (‘null’) mutants to wild-type (WT) flies with elevated oxidative stress levels induced by paraquat (PQ) feeding. A widely used herbicide, PQ is a well-studied drug often utilized experimentally to generate superoxide anions and induce oxidative stress in *Drosophila* (Parkes et al., 1993; Shukla et al., 2014). We examined synaptic transmission at the abdominal neuromuscular junction (NMJ) of 3^rd^ instar *Sod* larvae to WT larvae fed on PQ-containing medium from egg hatching. In adults, we compared the performance of the GF escape pathway (Engel & Wu, 1992) in *Sod* mutants to PQ-fed WT flies. The PQ feeding concentrations chosen for this study (5 or 10 mM throughout the larval stage; and 10 mM in adults for 4 d) were reported to induce lethality and behavioral abnormalities in WT flies within a 3 – 7 d window (Arking et al., 1991; Jahromi et al., 2013). Over this period, we indeed observed a marked loss of motor coordination. Flies often “stumbled” while walking and occasionally exhibited wing-buzzes upon gentle “tapping” of the vial. The PQ concentrations selected for exposing larvae throughout their development was near lethality levels, beyond which few larvae survived. Nevertheless, our neurophysiological findings reveal that PQ-fed WT individuals clearly do not replicate all *Sod* phenotypes in both larval and adult physiological recordings.

At the larval NMJ, we found *Sod* mutant larvae displayed a striking phenotype which was absent in WT counterparts (Figure 1A). Although an initial nerve stimulation produced similar excitatory junctional potentials (EJPs, analogous to EPSPs) in WT and *Sod* larvae (lower traces), after repetitive stimulation (at ~15 Hz, upper traces), we observed extra ‘humps’ in the *Sod* waveforms indicative of supernumerary EJPs (representative EJPs from a homozygous *Sod^n108^* and hetero-allelic *Sod^n108^/Sod^X39^* individuals are shown). These abnormal events indicate the stimuli recruit secondary action potentials in the motor nerve (Ueda & Wu, 2009) and suggest a form of nerve hyperexcitability in *Sod* mutants. In contrast, PQ-fed WT flies displayed EJPs that were remarkably similar to control diet-fed counterparts in terms of shape and amplitude. Indeed, across the population of larvae, extra ‘humps’ in the EJP waveform were commonly observed in *Sod* larvae but were exceedingly rare in PQ-fed WT or control-fed WT flies (Figure 1B).

**Figure 1.**
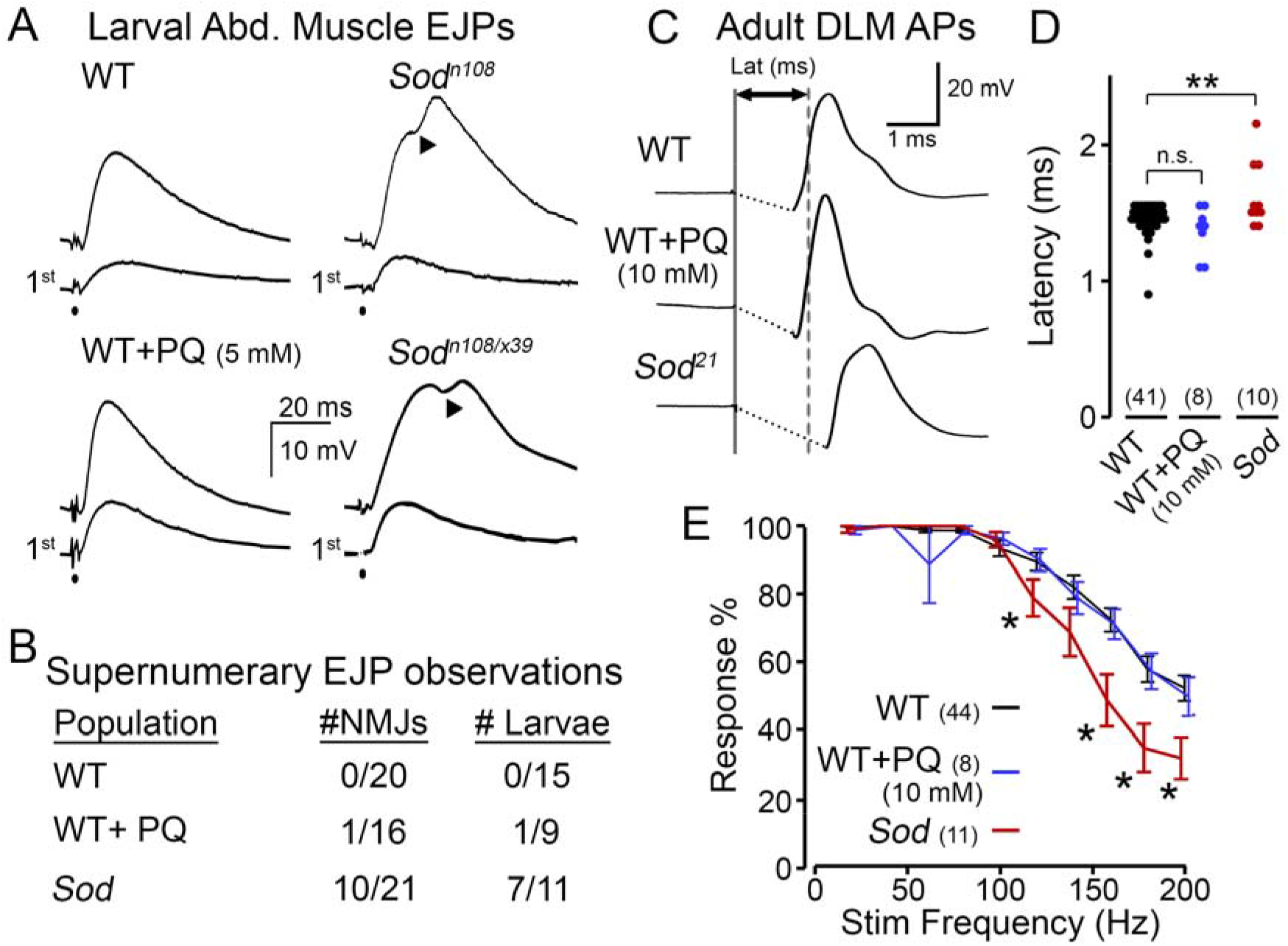
Comparison of *Sod* mutational and paraquat (PQ) pharmacological effects on electrophysiological properties of the *Drosophila* motor system. (A, B) Synaptic transmission at neuromuscular junctions (NMJs) of abdominal body-wall muscles in 3rd instar larvae. (A) Excitatory junctional potentials (EJPs) triggered by 0.1-ms motor axon stimulation. Upon repetitive nerve stimulation (15 Hz) supernumerary EJPs occur at NMJs in *Sod* mutants (arrowheads). The initial EJPs (“1st”) were not significantly different among WT control, PQ-fed WT, and *Sod* mutant larvae. However, subsequent nerve stimulation induced supernumerary EJPs (‘▲’ indicate humps in EJPs) in *Sod* larvae, but not in WT control or PQ-fed WT (see traces above the “1st” EJPs). (B) Proportion of NMJs and larvae displaying supernumerary EJPs. PQ concentration in food was 5 - 10 mM. Experiments were performed in HL3.1 saline (Feng et al., 2004) containing low levels of Ca^2+^ (0.1 - 0.2 mM). (C-E) Performance of the giant fiber (GF) jump-and-flight escape circuit in adult flies. (C) Electrical stimulation across the head directly activated GF pathway leading to flight muscle spikes monitored in dorsal longitudinal muscles (DLMs). The latency to spiking was measured as the duration between stimulus (solid line) and the DLM depolarization reaching its half-maximal amplitude (dashed line for WT). Stimulus artifacts have been removed for clarity. Note the retarded response in *Sod* mutant. (D) Scatterplot of the GF transmission latency for WT, PQ-fed WT and *Sod* mutants. Data points indicate latency from individual flies. (E) Frequency response of the GF pathway. Stimulus trains were delivered at progressively higher frequencies (20 – 200 Hz, see Methods), and the response rate was recorded. Note that *Sod* mutants display a relatively poor ability to follow high-frequency stimulation compared to PQ- and control-fed WT flies. For (D-E), number of flies studied are indicated in parenthesis.

To compare the effects of PQ feeding and *Sod* mutations on adult nervous system function, we examined stimulus-response relationships of the GF pathway. In this circuit, established visual and mechanosensory inputs trigger spiking in the GF neuron in response to visual looming or air pressure stimuli (Card & Dickinson, 2008; Lehnert et al., 2013; Mu et al., 2014; Trimarchi & Schneiderman, 1995). The descending GF axons project directly to the tergotrochanteral (TTM, ‘jump muscle’) motor unit and indirectly via the peripherally synapsing interneuron (PSI) to the dorsal longitudinal muscle (DLM, ‘flight muscle’) motor unit (Gorczyca & Hall, 1984; King & Wyman, 1980). Direct electrical stimulation of the GF neuron triggers an action potential in the DLM with a stereotypic latency in WT flies (~1.4 ms, Figure 1C), known as ‘short-latency’ responses (Engel & Wu, 1996). Consistent with previous observations (Iyengar et al., 2020), we found *Sod* flies displayed an overall increase in latency with a wider variability compared to WT counterparts (Figure 1C-D). However, PQ-fed WT flies did not show such retarded responses; instead latencies displayed were remarkably similar to control WT individuals. We also examined the ability of the GF pathway to follow trains of high-frequency stimulation (20 – 200 Hz) in PQ-fed flies and *Sod* mutants (Figure 1E). At stimulation frequencies higher than 100 Hz, we found *Sod* displayed a progressively worse ability to follow stimulation compared to WT counterparts (largely consistent with observations using a twin-pulse refractory protocol, c.f. Iyengar et al., 2020). However, across all frequencies, PQ-fed WT flies displayed a similar response rate compared to control diet-fed individuals.

Our observations at the larval NMJ and along the GF pathway in adults reveal clear and consistent defects in neurotransmission among *Sod* loss-of-function mutants. The *Sod* alleles we studied, i.e. *Sod^n108^*, *Sod^X39^* and *Sod^21^*, were independently isolated (Phillips et al., 1989; Şahin et al., 2017). Thus, it is most likely *Sod* disruption, rather than contributions from secondary ‘background’ mutations, gives rise to the phenotypes. We observed poor motor coordination and lethality, indicating our PQ-feeding indeed produced effects consistent with previous reports (Arking et al., 1991). Notably, even though the larval NMJ and adult GF pathway are remarkably robust systems, studies from our group and others have demonstrated clear electrophysiological alterations in *Sod* loss-of-function (this study; see also: Watson, et al., 2008, Iyengar, et al., 2020) In contrast, pharmacological manipulation of oxidative stress levels with standard PQ-feeding protocol failed to recapitulate *Sod* mutant phenotypes, suggesting the neuromuscular deficits in *Sod* flies do not arise by a simple overall elevation of superoxide levels. One possibility is that regulation of superoxide anions in specific cell types or subcellular compartments contribute to the complexity of *Sod* mutant phenotypes. Alternatively, mutations in *Sod* or PQ-exposure can induce different downstream adjustments of alterations, such as the level or activity of other redox enzymes (Missirlis et al., 2003) involving different 2^nd^ messenger signaling pathways (e.g. Nrf2 or NF-KB, Sies et al., 2017; Wang et al., 2018).

## Materials and Methods

### Fly Stocks and Drug Feeding

The *Sod* null mutant alleles examined included: *Sod^n108^*, *Sod^X39^* (Campbell et al., 1986; Phillips et al., 1989), and *Sod^21^* (Şahin et al., 2017). We studied homozygous *n108*, and *21* individuals; however, we examined the hetero-allelic *n108/x39* combination due to the poor viability of homozygous *x39* flies (Ruan & Wu, 2008). Phenotypes among the *Sod* alleles were remarkably similar. Therefore, the data was pooled for statistical analyses (Figure 1B, D & E). The WT strain was *Canton-Special*. For adult experiments, we studied WT and *Sod* populations before significant age-dependent mortality (< 1% and 5% mortality, 7-8 d for WT and 2-3 d *Sod*). For detailed characterization of the age-dependence of GF performance in *Sod* and WT flies, see Iyengar et al. (2020).

Flies were reared on Frankel & Brosseau (1968) cornmeal-based media (see Kasuya et al., 2019) at room temperature (23 °C). To make PQ-laced media, measured amounts of paraquat (Sigma #856177) and Blue #1 dye (to indicate mixing, flavorsandcolor.com, Diamond Bar, CA) were added to melted fly media to achieve final concentrations of 5 or 10 mM PQ and 25 μg/ml dye. For larval feeding experiments, larvae were reared on PQ media (5 mM or 10 mM) from egg hatching until the post-feeding 3^rd^ instar stage. Adults were aged to 3 – 4 d. and were then fed with PQ-containing (10 mM) medium for 4 d and used for recording.

### Larval Neuromuscular Electrophysiology

Neuromuscular recordings from post-feeding 3rd instar larvae are described in (Ueda & Wu, 2006, 2009). Dissections were performed in Ca^2+^-free HL3 saline (Stewart et al., 1994). Excitatory junction potentials (EJPs) were recorded from abdominal muscles in HL3.1 saline (Feng et al., 2004) containing relatively low Ca^2+^ levels (0.1 or 0.2 mM). To evoke nerve action potentials and EJPs, the segmental nerve was severed form the ventral ganglion and stimulated through the cut-end with a suction electrode (~10-μm inner diameter). The stimulation pulse train (0.1 ms stimuli delivered at 15 Hz for 50 s) was generated from a Grass S88 stimulator (model S88; Grass Instruments, West Warwick, RI) through a custom-made isolation unit. The stimulation voltage was adjusted to be double the nerve compound action potential threshold value to ensure motor axon stimulation. To record EJPs, a glass intracellular microelectrode filled with 3 M KCl (~ 60 MΩ resistance) was inserted in the abdominal muscle # 6/7 (Crossley, 1979). Signals were recorded by a DC amplifier (model M701, WPI, Sarasota, FL and an additional custom-built amplifier) and digitized using a data acquisition card (USB 6212, National Instruments, Austin TX) and pClamp5 (Molecular Devices, Burlingame, California, USA) controlled by a PC.

### Adult Giant-Fiber Physiology

Examination of the giant-fiber pathway physiology were performed in tethered flies (see Engel and Wu (1992); Iyengar and Wu (2014) for details). Flies were anesthetized on ice, affixed to a tungsten pin with cyanoacrylate glue, and given ~30 min to recover. To record spiking from the DLM, an electrolytically sharpened tungsten electrode was inserted into the attachment site of the dorsal-most fiber (#45, see Miller, 1950). A similarly constructed electrode was inserted into the abdomen as reference. Electrical activity was picked-up by an AC amplifier (gain: 100x, bandwidth: 10 – 10,000 Hz, AM systems Model 1800) and digitized by data acquisition card (USB 6212, National Instruments) controlled by a PC running LabVIEW (2018, National Instruments).

Electrical stimulation to activate the GF pathway was delivered by an isolated pulse stimulator (AM systems Model 2100) via sharpened tungsten electrodes inserted in each cornea. Stimuli consisted of brief pulses (0.1 ms duration, 30 V amplitude) which reliably activated the GF neuron and downstream elements (‘short-latency responses’ in Engel & Wu, 1996; Iyengar et al., 2020). The GF latency (Figure 1D) was determined as the interval between stimulation and the DLM spike reaching its half-maximum height. The frequency following ability of the GF pathway was determined by delivering trains of 10 pulses at increasing frequencies (20 Hz – 200 Hz in 20 Hz increments) with a 5-s interval between trains. For each pulse train, the number of DLM responses divided by number of stimuli was reported.

### Statistical Analysis

All statistical analysis was done in MATLAB (r2020b, MathWorks). Kruskal-Wallis non-parametric ANOVA tests (with Holm-Bonferroni corrected rank-sum *post hoc* analysis) were employed to establish statistical significance in Figure 1D and E. * p < 0.05, and ** p < 0.01.

## Acknowledgments

We thank the undergraduate research assistants in the Wu Lab for their help in maintaining fly stocks and assisting with experiments. This work was supported by an NIH Grants (AG 051513 and NS 111122 to CFW). AI was supported by an Iowa Neuroscience Institute Fellowship.

